# Task-based value generalization correlates with positive overgeneralization and bipolar traits

**DOI:** 10.64898/2026.07.12.737635

**Authors:** Jing Li, Maya Malaviya, Daniel Bennett, Angela Radulescu

**Author notes:** Corresponding authors: Jing Li and Angela Radulescu.

## Abstract

Positive overgeneralization – the tendency to generalize from specific successes to broad expectations of future reward – has been linked to vulnerability to mania. Because positive overgeneralization has only been assessed using self-report measures, we have limited insight into the underlying cognitive process. Here, we introduce a behavioral paradigm that quantifies how reward generalizes to novel stimuli, and characterize individual generalization profiles by fitting psychometric functions to choice data. In an online transdiagnostic study (*N* = 163), broader task-based generalization was associated with both higher self-reported positive overgeneralization and subclinical bipolar traits. Response times provided converging evidence: a smaller increase in response time as stimuli became less similar to the rewarded stimulus was associated with higher positive overgeneralization. In computational simulations, we show that self-efficacy, operationalized as the relative weight placed on the best and worst available future actions, shapes generalization, with more weight on the best possible action broadening the spread of expected reward to similar states. Together, these findings show that positive overgeneralization can be measured behaviorally, and identify one computational factor that may determine how broadly reward generalizes.

## 1 Introduction

Success can dramatically reshape motivation, sometimes fueling escalated confidence and increased goal pursuit. In its extreme form, this pattern is characteristic of mania, a hallmark of bipolar disorder marked by episodic surges in energy, elevated mood, and expansive goal-directed behavior. Hypomania involves a milder form of the same symptom profile that does not reach the threshold for a full manic episode (American Psychiatric Association, 2013).

This clinical phenotype has motivated a substantial literature linking mania vulnerability to dysregulation of the Behavioral Activation System (BAS), a neurobehavioral system governing approach motivation and sensitivity to reward cues (Johnson, Edge, Holmes, & Carver, 2012). BAS sensitivity, commonly assessed using the Behavioral Inhibition/Behavioral Activation Scales (BIS/BAS) (Carver & White, 1994), prospectively predicts both the onset of hypomanic episodes and the severity of manic symptoms over time (Alloy et al., 2008). Consistent with this heightened approach motivation, individuals at risk for bipolar disorder also demonstrate increased willingness to expend effort toward difficult-to-obtain goals (Johnson & Carver, 2006). Hypomanic traits have further been linked to overly positive cognitive biases following success, including self-serving attributions that even extend to chance-based outcomes, where personal ability plays no objective role (T. D. Meyer, Barton, Baur, & Jordan, 2010). However, the cognitive processes that sustain increased behavioral activation following initial success remain unclear.

One cognitive mechanism through which BAS dysregulation may operate is positive overgeneralization (POG) — the tendency to respond to a specific success with excessive confidence and to generalize from that positive experience to broad expectations about the self and future outcomes (Eisner, Johnson, & Carver, 2008; Stange et al., 2012). From a reinforcement learning perspective, POG can be conceptualized as an alteration in how reward value propagates through a state space (Gershman & Niv, 2015; Radulescu & Niv, 2019; Sutton & Barto, 2018; Wimmer, Daw, & Shohamy, 2012; Wu, Meder, & Schulz, 2025). Rather than assigning increased expected value only to the specific actions or contexts that led to success, individuals high in POG appear to assign value to loosely related states, goals, or self-beliefs.

For example, solving a difficult technical problem at work may not only increase the value of related professional behaviors, such as taking on more ambitious projects, but also inflate expectations of success in unrelated life domains, such as romantic relationships or athletic performance. Importantly, upward POG prospectively predicts increases in hypomanic symptoms, particularly among individuals high in BAS sensitivity (Stange et al., 2012).

Despite its relevance to (hypo)mania, the construct of POG has been developed largely within the domain of self-report, most notably through the Positive Generalization Scale (Eisner et al., 2008). While self-report provides valuable insight into individuals’ subjective beliefs about their generalization tendencies, it does not reveal the processes that give rise to those tendencies. In the context of positive experiences, a key question is how information about reward is generalized beyond the experience itself to shape expectations in new situations. This process could reflect differences in value updating, the breadth of generalization, or credit assignment. Characterizing this process would enable formal quantification of POG, adjudication between mechanistic accounts, and more direct links to underlying neural substrates.

To address this gap, we introduce a reward generalization paradigm designed to quantify individual differences in how broadly learned value extends to novel, but similar, stimuli. Participants first learn the reward value of distinct stimulus shapes and then make reward predictions for intermediate, untrained stimuli. We characterize each participant’s generalization profile by fitting a psychometric function to reward predictions, yielding a parameter that indexes the breadth of value generalization. We hypothesized that broader behavioral generalization would be associated with higher self-reported (1) positive overgeneralization and (2) bipolar spectrum traits. To investigate what computational mechanism might produce differences in generalization breadth, we implemented a reinforcement learning model in which self-efficacy — an agent’s belief about its ability to attain the best available future outcome — modulates the influence of anticipated future value during learning (Li & Radulescu, 2024; Zorowitz, Momennejad, & Daw, 2020).

Across an online transdiagnostic sample, we find that two task-based measures — the breadth of value generalization and the rate at which response time changes with distance from a participant’s own choice boundary — both track self-reported positive overgeneralization. Generalization breadth also relates to subclinical bipolar traits, while neither measure relates to current depressive or anxiety symptoms. Exploratory analyses suggest that the response-time slope may track hypomanic rather than depressive traits, although this pattern did not survive correction for multiple comparisons. Simulations show that varying the self-efficacy parameter is sufficient to shift the breadth of value propagation.

## 2 Methods

### 2.1 Overview

Participants completed a single online session that lasted approximately 30 mins, consisting of an instruction phase, training, test, and a questionnaire battery (Fig. 1A). The session consisted of a behavioral task (“Fossil Hunt”), followed by a battery of self-report questionnaires. The Fossil Hunt task included an instruction phase with comprehension checks, a training phase, and a test phase. The study was implemented in PsychoPy (Peirce et al., 2019) and hosted on Pavlovia (https://pavlovia.org). Participants were recruited via Prolific and compensated at a rate of $12/hour, with up to $4 in performance-based bonuses.

**Figure 1:**
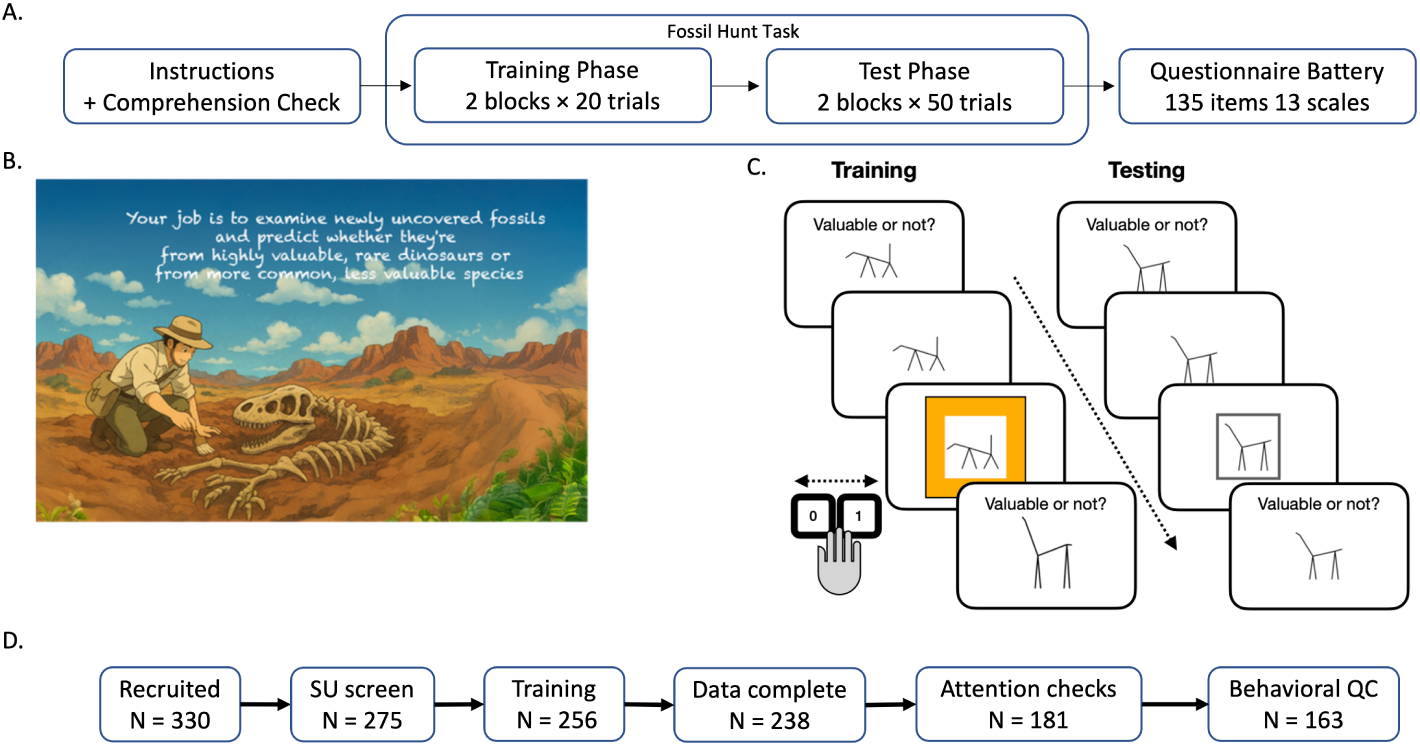
Study design. **A:** Session timeline. Participants completed a single online session consisting of the Fossil Hunt behavioral task followed by a battery of self-report questionnaires. **B:** Fossil Hunt game introduction screen. Participants played the role of fossil hunters learning to identify valuable dinosaur specimens. **C:** Task structure. During training, participants learned to associate one dinosaur skeleton shape with reward (“valuable”) and received feedback on each trial. During the test phase, participants judged whether novel stimuli with shapes intermediate between the two training exemplars would be valuable, using a binary keypress (0/1). **D:** Participant retention. Sequential application of substance use screening, training performance, data completeness, survey attention check, and behavioral quality control criteria yielded a final analytic sample of *N* = 163.

### 2.2 Participants

A total of *N* = 330 participants were recruited online via Prolific. Exclusion proceeded in five sequential stages (Fig. 1D).

First, participants completed a substance use screening prior to the task. Participants were asked whether their use of drugs or medications in the past year had caused functional impairment (e.g., problems with family or others, worsened health, reduced activities, withdrawal symptoms, or use in physically risky situations). To minimize confounds from substance-related alterations in reward processing, those endorsing substance-related impairment were prevented from proceeding with the full experiment (*n* = 55 excluded; *N* = 275 remaining).

Second, participants who failed to perform above chance during the training phase were excluded (*n* = 19 excluded; *N* = 256 remaining), ensuring that remaining participants had successfully learned the reward contingencies before proceeding to the test phase.

Third, participants whose data were partially lost due to server errors during the session were excluded (*n* = 18 excluded; *N* = 238 remaining).

Fourth, seven embedded attention-check items with known correct responses were distributed across the self-report questionnaire battery (e.g., “To answer this question, please select option [X]”); participants who answered any item incorrectly were excluded (*n* = 57 excluded; *N* = 181 remaining). This screening step was motivated by evidence that careless or inattentive survey responding can induce spurious associations between task behavior and self-report symptom measures, particularly when symptom endorsement is positively skewed in the sampled population (Zorowitz, Solis, Niv, & Bennett, 2023).

Fifth, behavioral quality-control criteria were applied to test-phase data (*n* = 18 excluded; *N* = 163 remaining). Prespecified criteria assessed monotonicity of the generalization gradient, response times, data completeness, and response bias. The monotonicity criterion ensured that participants treated stimuli more similar to the rewarded shape as more valuable — the basic signature of having learned the reward contingency and applied it along the similarity continuum. Twelve participants were excluded for failing to show a monotonically increasing relationship between stimulus shape and proportion “valuable” responses (Spearman *ρ < ρ*_crit_ = .55, one-sided *α* = .05), including three with inverted gradients suggesting confusion of the reward contingencies. Six additional participants were excluded for excessive slow responding (*>*20% of trials exceeding 3s).

No participants were excluded for responding too quickly or insufficient data. The final analytic sample consisted of *N* = 163 participants.

### 2.3 Behavioral task

We developed “Fossil Hunt”, a gamified behavioral task in which human participants played the role of paleontologists learning to identify valuable dinosaur specimens (Fig. 1B). Our goal was to induce learning of a value gradient along a single, initially value-neutral stimulus dimension by training participants on two maximally dissimilar extremes, one associated with reward and the other not. We then asked how participants generalize learning from the two extremes to the intermediary stimuli.

#### 2.3.1 Stimuli

Stimuli consisted of images of stick figures representing quadrupeds, adapted from previous work on perception and categorization (Malaviya, Sucholutsky, Oktar, & Griffiths, 2022; Olman & Kersten, 2004; Sanborn & Griffiths, 2007). Each figure was defined by 9 continuous features describing the configuration of its body and limbs. We selected two endpoint figures and generated eight intermediate stimuli by interpolating between their feature values in evenly spaced steps. Because all 9 features were interpolated together, the resulting stimuli varied along a single dimension — the position of each shape between the two endpoints. A separate similarity-rating study confirmed that they were also evenly spaced perceptually (Supplementary Section S2). Throughout the experiment, the stick figure quadrupeds were referred to as “dinosaur fossils”.

#### 2.3.2 Task timeline

On each trial of the training phase, participants viewed one of two endpoint fossils — the shapes at either extreme of the stimulus continuum (Fig. 1C; see Fig. S1 for the full stimulus set). One of the endpoints (counterbalanced across participants) was associated with reward, while the other was not. Reward contingencies were probabilistic: the rewarded shape yielded reward on 85% of trials, while the non-rewarded shape yielded reward on 15% of trials. Training consisted of two training blocks of 20 trials each (40 trials total).

During the test phase, participants were presented with eight novel stimuli that interpolated between the two training exemplars (Fig. 1C). On each trial, participants saw one stimulus and predicted whether it would lead to reward by pressing one of two keys (0 = not valuable, 1 = valuable). They were instructed to base their judgments on each stimulus’s similarity to the shapes they had learned during training. To prevent further learning and isolate generalization, participants received no feedback. Only participants who performed above chance during training were retained, so all participants in the analytic sample had learned the reward contingencies before the test phase (see Participants).

The test phase consisted of two blocks, each presenting the full set of 10 stimulus shapes (eight novel intermediate stimuli plus the two trained exemplars). Participants completed 100 test trials (10 per stimulus shape) across the two blocks. Before each test block, participants answered forced-choice comprehension questions assessing understanding of the response keys, bonus structure, and the relationship between stimulus similarity and value. Participants were required to answer all questions correctly before proceeding; incorrect responses triggered a loop back to the instructions.

Participants could earn up to $4 in performance-based bonuses contingent on accuracy during both training and test phases. To incentivize engagement without biasing generalization responses, only aggregate bonus earnings were revealed at the end of each block. Correctness feedback was never provided for individual trials.

### 2.4 Self-report measures

Following the behavioral task, participants completed 12 self-report instruments assessing traits and symptoms relevant to bipolar disorder (Table 1). The Positive Overgeneralization (POG) scale (Eisner et al., 2008) yields a total score and three subscales: upward generalization (expecting success at more ambitious goals within the same domain), lateral generalization (expecting success in other domains), and social generalization (expecting broader social success after a positive social experience). The 7Up/7Down composite sums all 14 items of the 7 Up 7 Down Inventory, which is conventionally scored as two scales; 7-Up and 7-Down were also scored separately. The administered BIS/BAS subset comprised 12 items, yielding three four-item scores (BIS, BAS Drive, and BAS Reward Responsiveness); BAS Fun Seeking was not included. Seven attention-check items were embedded across the battery, each with a known correct response (e.g., “To answer this question, please select option [X]”), following the approach of Zorowitz et al. (2023). Attention-check items were excluded from all questionnaire scores and item-level analyses, and participants who answered any check incorrectly were excluded (see Participants).

**Table 1:** Self-report measures, grouped by their role in the bipolar-spectrum hypothesis. The 12 instruments yield 19 questionnaire scores: POG total and its three subscales (upward, lateral, social); the 7Up/7Down composite, 7-Up, and 7-Down; BIS, BAS Drive, and BAS Reward Responsiveness; and one score from each of the other nine instruments.

| Measure | Construct | Reference |
| --- | --- | --- |
| *Bipolar spectrum and mania risk measures* |  |  |
| POG | Positive overgeneralization (total and 3 subscales) | Eisner et al. (2008) |
| 7Up/7Down | Subclinical bipolar traits (total and 2 subscales) | Youngstrom et al. (2013) |
| PMQ-9 | Current manic symptoms | Cerimele et al. (2022) |
| PHQ-9 | Current depressive symptoms | Kroenke et al. (2001) |
| BIS/BAS | Behavioral inhibition / activation (3 subscales) | Carver & White (1994) |
| WASSUP | Ambitious goal-setting | Johnson & Carver (2006) |
| NGS | Narcissistic grandiosity | Rosenthal et al. (2020) |
| GSE | General self-efficacy | Schwarzer & Jerusalem (1995) |
| *Additional measures* |  |  |
| GAD-7 | Current anxiety symptoms | Spitzer et al. (2006) |
| PSWQ | Worry | Meyer et al. (1990) |
| SHAPS | Anhedonia | Snaith et al. (1995) |
| IE-4 | Locus of control | Nießen et al. (2022) |

### 2.5 Behavioral modeling

We modeled each participant’s reward predictions with a four-parameter logistic psychometric function. As novel stimuli become increasingly similar to a previously rewarded exemplar, the probability of predicting reward should increase in a graded fashion, and a sigmoidal function captures this transition on a bounded probability scale. Because reward assignment to the two end-point shapes was counterbalanced, all analyses used an axis defined by distance from each participant’s own rewarded exemplar: *d* = 0 at the trained rewarded endpoint and *d* = 90 at the trained non-rewarded endpoint. Predictions were modeled as

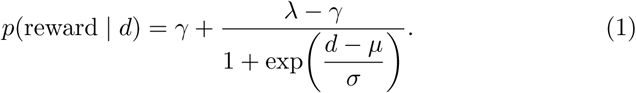

The logistic form (Wichmann & Hill, 2001) allows four parameters to vary across individuals (Table 2). Larger *µ* values indicate that the transition to predicting stimuli as “valuable” occurs farther from the trained rewarded shape and therefore denote broader reward generalization; at *d* = *µ*, predicted reward probability equals the midpoint between the two asymptotes, (*γ* + *λ*)*/*2. Parameters were estimated via nonlinear least-squares using Python’s scipy optimization library, and all four were recoverable in simulation (*r >* .91; Supplementary Sections S3 and S4).

**Table 2:** Parameters of the four-parameter logistic psychometric function.

| Parameter | Interpretation | Analysis |
| --- | --- | --- |
| $\mu$ | Distance from the rewarded exemplar at which reward predictions reach their midpoint | Primary |
| $\sigma$ | Width of the transition | Secondary, via $\log S$ |
| $\gamma$ | Reward prediction rate at the nonrewarded end | Descriptive |
| $\lambda$ | Reward prediction rate at the rewarded end | Descriptive |

We take *µ* as our primary index of generalization breadth, since it locates the transition along the stimulus continuum. Because *σ* describes the width of that transition rather than its slope, we index the sharpness of the boundary with the maximum derivative *S* = (*λ − γ*)*/*(4*σ*) and analyze log *S* in secondary analyses (Supplementary Section S7). The asymptotes index response tendencies at the trained endpoints rather than generalization, and are reported descriptively, as is the area under the fitted curve (AUC), a summary of overall “valuable” responding.

### 2.6 Response time analysis

Response times (RTs) from test trials were analyzed as a function of perceptual distance from each participant’s fitted psychometric boundary (i.e. the subjective midpoint at which the participant switches their prediction). RTs outside 0.2–8.0 s were excluded (0.7% of trials), and retained values were natural-log transformed. Raw RTs were strongly right-skewed (skewness = 3.57); the natural-log transformation reduced this to 1.04 and was used in all models (Fig. S8).

For each participant, we regressed log RT on distance from their fitted boundary. Distance was expressed in 10-unit steps — the spacing between adjacent stimuli — and signed so that positive values point toward the rewarded endpoint,

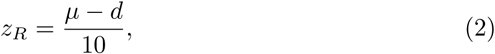

and log RT was modeled as

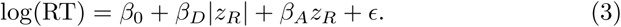

The model has three coefficients (Table 3). The intercept *β*_0_ is log RT at the choice boundary; the absolute-distance term *β_D_* captures how RT changes as stimuli move away from the boundary in either direction; and the signed term *β_A_* captures any difference between the two directions.

**Table 3:** Coefficients of the response time analysis.

| Coefficient | Interpretation |
| --- | --- |
| $\beta_0$ | Log RT at the choice boundary |
| $\beta_D$ | Change in log RT per stimulus step away from the boundary |
| $\beta_A$ | Difference between the two directions |

Correlations with self-report used each participant’s fitted coefficients. Because distance from the boundary is computed from a fitted parameter rather than a known quantity, we verified through simulation that both coefficients could be recovered when error in the boundary estimate was propagated into the RT model (Supplementary Section S4).

### 2.7 Exploratory factor analysis

To identify additional latent dimensions of self-report, we conducted exploratory factor analyses on all 135 substantive questionnaire items from the 12-instrument battery, excluding the seven attention checks. Items were organized into 13 groups for loading displays and purity calculations, with 7-Up and 7-Down represented separately. Because the appropriate number of factors to retain in exploratory factor analysis has no single agreed-upon criterion, it is common practice to evaluate multiple resolutions (Costello & Osborne, 2005). Consistent with this, in our data, parallel analysis (Horn, 1965) indicated that *k* = 9 factors should be retained, while the scree elbow suggested *k* = 3 (Fig. S6). To illustrate how associations with *µ* differ as a function of resolution (i.e. number of factors), we report results across all factor solutions from *k* = 2 to *k* = 12. We apply FDR correction separately at each *k*, across the *k* factor–*µ* correlations extracted at that resolution (e.g., four comparisons at *k* = 4, nine comparisons at *k* = 9).

To characterize the contribution of different self-report scales to each factor, we computed a descriptive metric we term “scale purity,” which quantifies the proportion of a factor’s total squared item loadings attributable to items from a given scale. For a target factor *f* and a self-report scale *X*, scale purity is defined as:

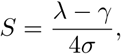

Here, *λ_i,f_* is the loading of item *i* on factor *f*, and the denominator sums over all 135 items in the self-report data *Y*. Items were assigned to scales based on their membership in the original self-report instruments (e.g., all 16 POG items contributed to the POG scale group). Purity values range from 0 (no contribution from scale *X*) to 1 (factor entirely defined by items from scale *X*). Items were grouped by their source instrument, so instruments with subscales (i.e., POG, BIS/BAS, 7Up/7Down) form a single group for this analysis.

### 2.8 Statistical analyses

We tested two primary hypotheses: that broader reward generalization (larger *µ*) would be associated with higher (1) POG total score and (2) 7Up/7Down composite score. Both were tested using two-sided, unadjusted *p*-values with *α* = .05. The hypotheses and exclusion criteria were specified in advance of analysis, although they were not formally preregistered.

We also report the full profile of associations between each behavioral measure and all 19 questionnaire scores, with 7-Up and 7-Down entered separately to assess hypomanic and depressive tendencies. For *µ*, POG total and the 7Up/7Down composite were the primary tests and are reported unadjusted; multiple comparisons were controlled using the false discovery rate (FDR) across the remaining 17 scores (the three POG subscales, 7-Up, 7-Down, the three BIS/BAS scores, and the nine other questionnaire scores). The log-RT distance slope *β_D_*, log psychometric steepness, and the directional RT measures *β_A_* and *β*^+^ were analyzed after the primary *µ* tests and are secondary; for each of these measures, FDR correction was applied separately across all 19 scores. Exploratory factor-level associations were corrected within each factor solution across the *k* factor–*µ* correlations. All correlation tests were two-sided, and *q* denotes a FDR-adjusted *p*-value.

## 3 Results

### 3.1 Psychometric characterization

Across the test phase, participants judged intermediate stimuli as more or less valuable depending on their similarity to previously rewarded shapes. Behavior in this phase revealed substantial individual differences in generalization (Fig. 2A). Figure 2B illustrates three representative participants: one with an intermediate generalization profile (*µ* = 48, near the midpoint of the stimulus range at 45), one generalizing broadly (*µ* = 68), and one generalizing narrowly (*µ* = 32). These examples show that larger *µ* reflects a greater tendency to judge stimuli farther from the rewarded exemplar as valuable.

**Figure 2:**
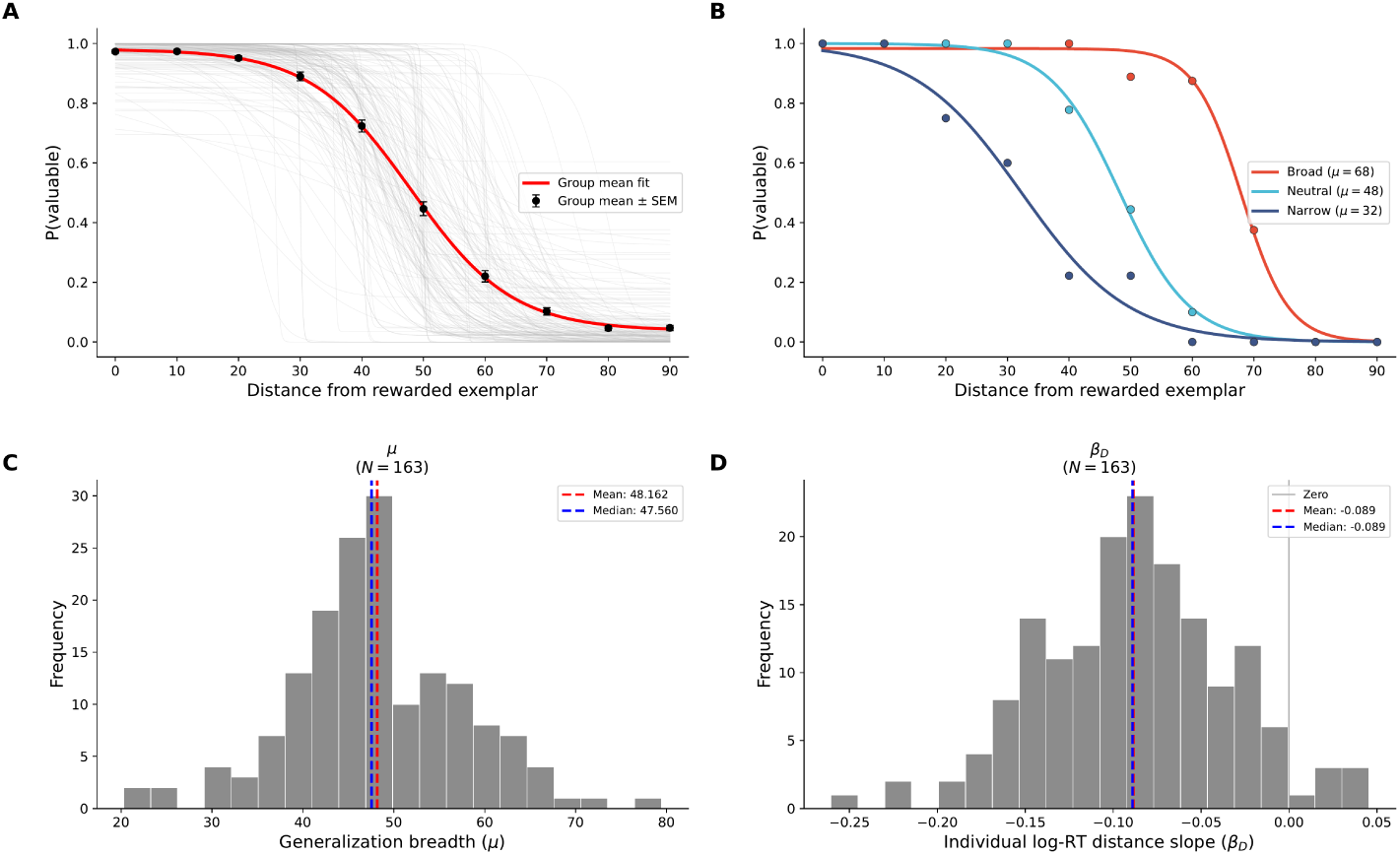
Generalization behavior and model fits. **A:** Group-level generalization curve. Black points show mean proportion “valuable” responses (*±* SEM) at each distance from the rewarded exemplar; the red curve shows the four-parameter logistic fit to group-averaged data (*µ* = 48.00, *σ* = 8.20, *γ* = .038, *λ* = .981); gray lines show individual fits. **B:** Three example participants with broad (*µ* = 68), intermediate (*µ* = 48), and narrow (*µ* = 32) generalization profiles; points show observed proportions, curves show fitted functions. **C:** Distribution of *µ* (*M* = 48.16, *SD* = 9.48). **D:** Distribution of the log-RT distance slope *β_D_* (*M* = *−*.089, *SD* = .053). Dashed red and blue lines in C and D indicate the mean and median. *N* = 163.

#### 3.1.1 Parameter distributions

The fitted midpoint *µ* showed wide individual variability (*M* = 48.16, *SD* = 9.48, range: 20.29–79.45; Fig. 2C), indicating large individual differences in generalization breadth across participants. The transition scale *σ* was right-skewed (*M* = 5.07, *SD* = 4.15; Fig. S4A), indicating that most participants switched between “valuable” and “not valuable” over a narrow range of stimuli, with a subset transitioning almost abruptly. Psychometric steepness was examined separately as a secondary measure (Supplementary Section S7). The lower asymptote *γ* was near floor for most participants (*M* = .05, *SD* = .09; Fig. S4C), and the upper asymptote *λ* near ceiling (*M* = .97, *SD* = .05; Fig. S4B), indicating that most participants reliably discriminated the trained exemplar stimuli. These values indicate that participants who passed the training check responded near-perfectly at the trained shapes, confirming that the reward contingency was learned. Because the endpoints were highly discriminable, the asymptotes showed little variation across participants and were not analyzed further as individual differences. The fitted parameters were largely independent of one another, with the exception of AUC, which overlaps with *µ* by construction and is therefore reported descriptively (Supplementary Section S5).

### 3.2 Response time characterization

Response times were fastest at the two trained endpoints and rose toward each participant’s fitted choice boundary (Fig. 3). Median RT was longest near the middle of the stimulus range (.963 s at *d* = 40 and 1.017 s at *d* = 50) and shortest at the two trained endpoints (.667 s and .730 s). RT peaked near the boundary rather than changing steadily across the stimulus range.

**Figure 3:**
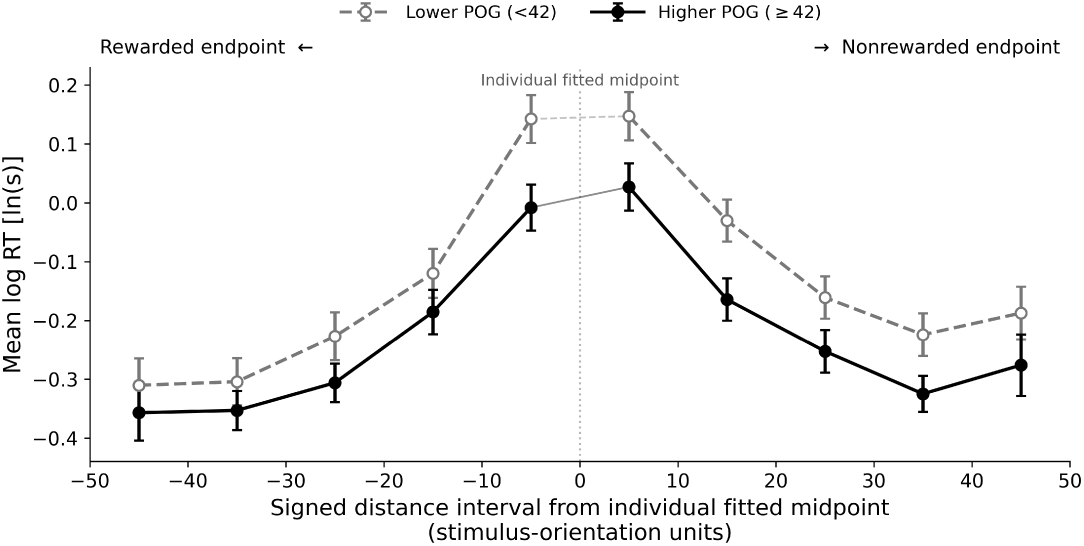
Response times by distance from the choice boundary. Mean log RT by signed distance from each participant’s fitted midpoint, split at the median POG total score (POG = 42). Negative values indicate stimuli between the midpoint and the rewarded endpoint. Trials were averaged within participant and bin before averaging across participants; error bars are SEM across participant means. The median split is for display only; all reported statistics use continuous POG scores. *N* = 163.

The mean distance slope was *β_D_* = *−*.089 (*SD* = .053; Fig. 2D), reliably below zero, *t*(162) = *−*21.37, *p <* .001, confirming that RT tracked distance from the boundary for most participants. Responses also slowed asymmetrically: the mean directional coefficient *β_A_* = *−*.006 (*SD* = .028) differed from zero, *t*(162) = *−*2.73, *p* = .007, corresponding to slopes of *−*.095 toward the rewarded endpoint and *−*.082 toward the nonrewarded endpoint.

Generalization breadth and the distance slope were uncorrelated (*r* = .03, *p* = .732; Supplementary Section S5), so the two measures were treated as separate correlates of self-report in the analyses that follow.

### 3.3 Relationship between reward generalization and self-report

In confirmatory analyses, we tested whether broader task-based generalization is associated with higher self-reported POG. Building on prior work linking POG to bipolar risk (Johnson & Carver, 2006; Stange et al., 2012), we further predicted that value generalization would also be associated with bipolar traits.

#### 3.3.1 Positive overgeneralization

Consistent with our first hypothesis, *µ* showed a significant positive association with total score on the POG questionnaire (Fig. 4A). Participants with higher POG scores exhibited larger *µ* values, indicating broader generalization of “valuable” judgments to stimuli farther from the trained rewarded exemplar (Pearson *r* = .21, *p* = .008, *N* = 163). This relationship also held for key subscales, with the strongest effect for the upward generalization dimension (*r* = .23, *p* = .003), followed by the social dimension (*r* = .19, *p* = .015). In contrast, the lateral subscale was not significantly associated with *µ* (*r* = .12, *p* = .121). These results indicate that individuals who overgeneralize positive outcomes to elevated self-aspirations and social success also exhibit broader task-based reward generalization.

**Figure 4:**
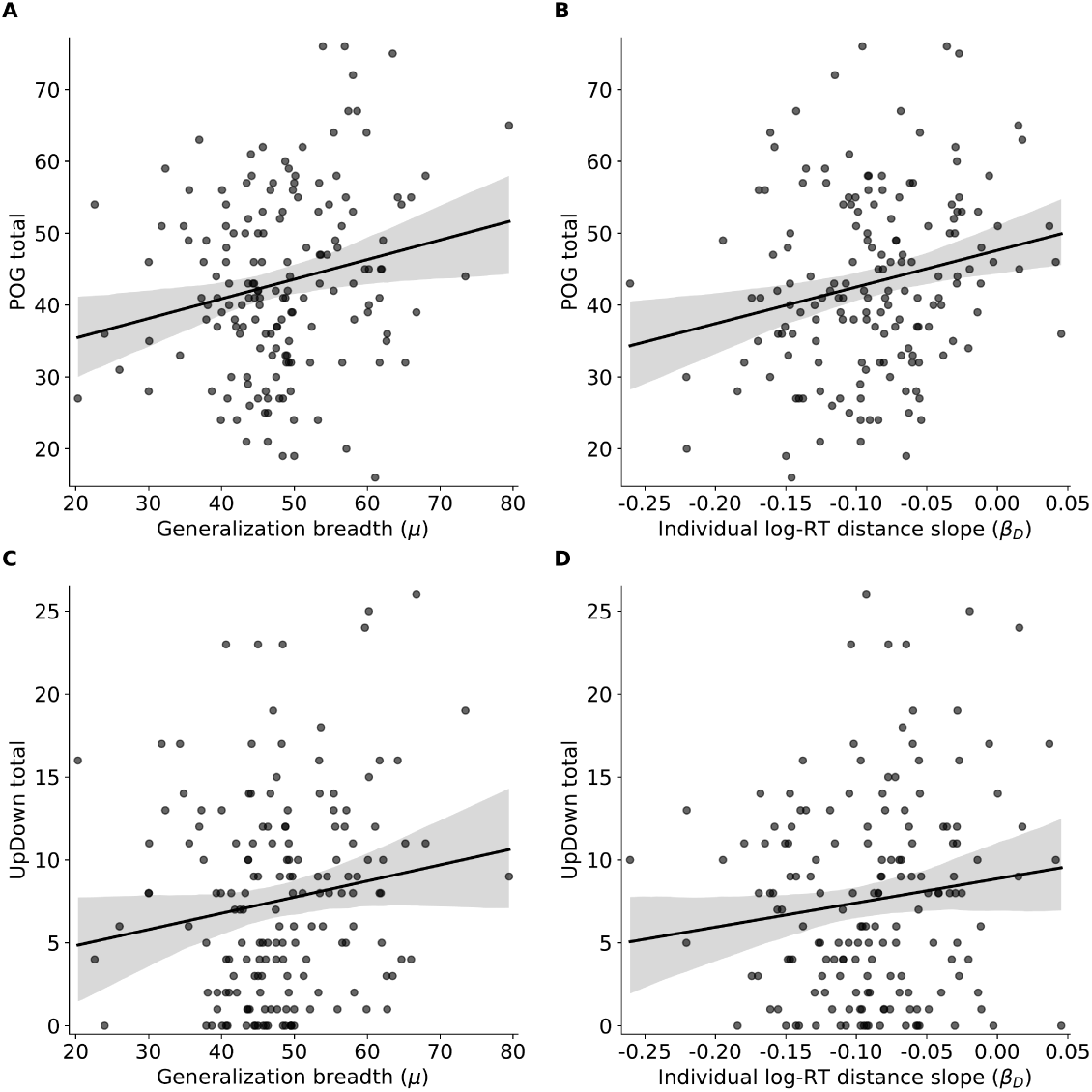
Task measures against self-report. **A:** Generalization breadth *µ* against POG total score. **B:** Log-RT distance slope *β_D_* against POG total score. **C:** *µ* against 7Up/7Down total score. **D:** *β_D_* against 7Up/7Down total score. Black lines show least-squares fits with 95% confidence bands. *N* = 163.

The secondary response time measure converged in the same direction, such that higher POG was associated with a shallower rise in response time as stimuli moved from the rewarded shape toward each participant’s own choice boundary (Fig. 4B; *r* = .22, *p* = .005).

#### 3.3.2 Bipolar spectrum traits

Consistent with our second hypothesis, *µ* was positively associated with the 7Up/7Down total score (Fig. 4C; *r* = .15, *p* = .050), such that greater bipolar-spectrum trait scores corresponded to broader generalization. The two subscales of the inventory behaved differently. Generalization breadth tracked 7-Down, which indexes trait-like depressive tendencies (*r* = .16, *p* = .041, *q* = .232), but not 7-Up, which indexes hypomanic tendencies (*r* = .06, *p* = .426, *q* = .757). For the secondary log-RT distance slope, the pattern was the opposite: the slope tracked 7-Up (*r* = .18, *p* = .022, *q* = .084) but not 7-Down (*r* = .03, *p* = .682, *q* = .807), and showed no association with the composite (Fig. 4D; *r*= .13, *p* = .102, *q* = .242). The associations with the subscales did not survive FDR correction, so we report the reversal as an exploratory descriptive pattern rather than a demonstrated dissociation.

Across the full 19-score profile, the largest correlation for each measure was with POG score (POG upward for *µ*, *r* = .23; POG total for *β_D_*, *r* = .22; Fig. 5). The largest remaining association was *β_D_* with BAS Drive (*r* = .19, *p*= .018); neither this nor any other survived FDR correction (Table 4). Both measures showed near-zero associations with current depressive (PHQ-9) and anxiety (GAD-7, PSWQ) symptoms.

**Figure 5:**
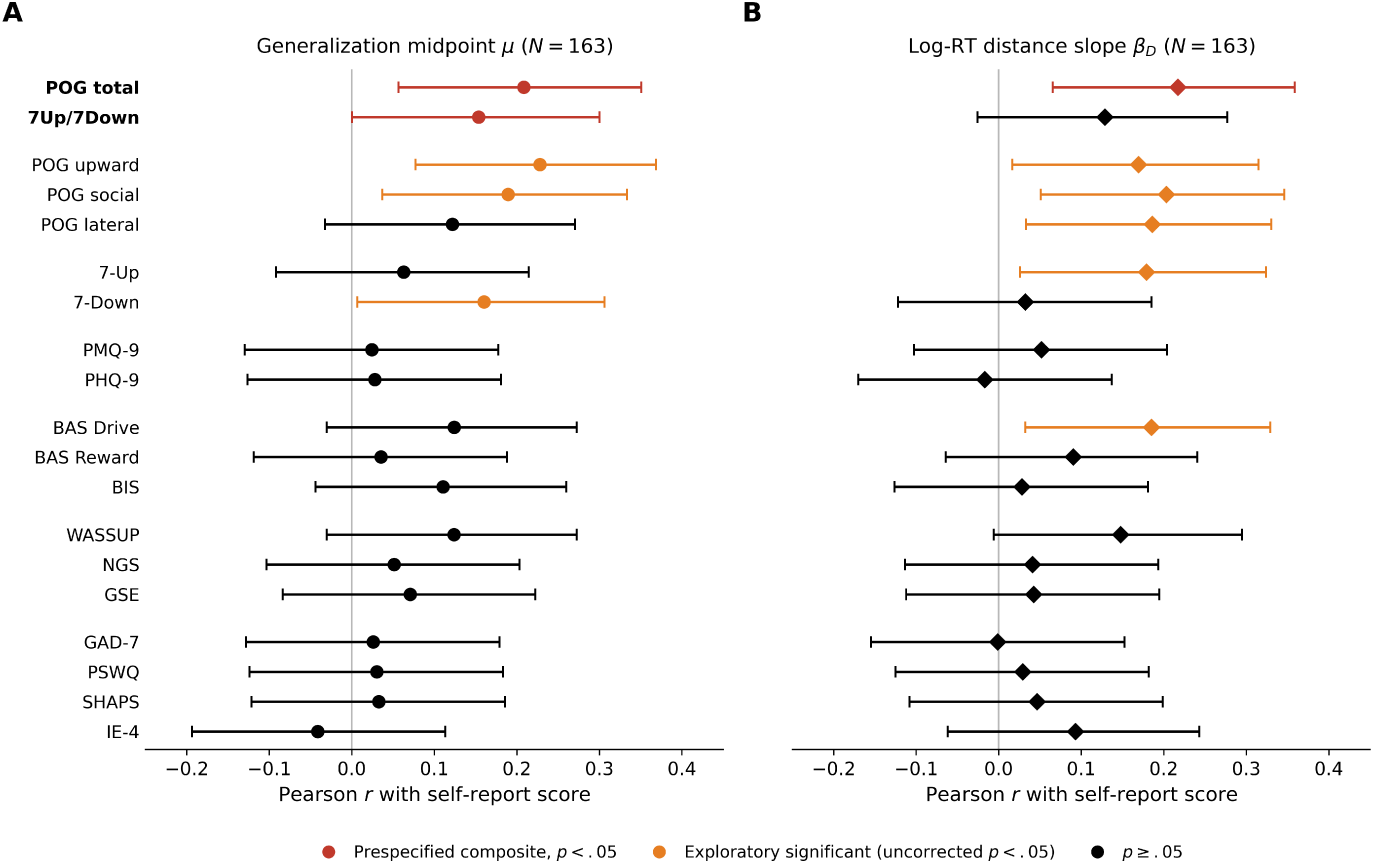
Self-report correlates of *µ* and *β_D_*. Pearson correlations with 95% confidence intervals (error bars) between each self-report score and **(A)** the generalization midpoint *µ* and **(B)** the log-RT distance slope *β_D_* (*N* = 163). Bold marks the two total scores whose *p*-values were not corrected: POG total and the 7Up/7Down composite. The remaining 17 scores were FDR-corrected within each panel. Red marks a confirmatory hypothesis test with *p < .*05; orange marks an exploratory significant association (any other score with uncorrected *p < .*05; none survived FDR); black marks *p ≥ .05*. Raw *p*-values and corrected *q*-values are reported in Table 4.

**Table 4:** Pearson correlations of all 19 questionnaire scores with generalization breadth *µ* and the log-RT distance slope *β_D_* (*N* = 163). For *µ*, POG total and the 7Up/7Down composite were the primary tests and are reported unadjusted (*q*: –); FDR was applied across the remaining 17 scores. For *β_D_*, a secondary measure, no score was exempt and FDR was applied across all 19 scores. All *p*-values are two-sided and unadjusted.

| Score | $\mu$ | | | $\beta_D$ | | |
| --- | --- | --- | --- | --- | --- | --- |
| | $r$ | $p$ | $q$ | $r$ | $p$ | $q$ |
| POG total | .21 | .008 | – | .22 | .005 | .084 |
| 7Up/7Down | .15 | .050 | – | .13 | .102 | .242 |
| POG upward | .23 | .003 | .058 | .17 | .031 | .097 |
| POG social | .19 | .015 | .131 | .20 | .009 | .084 |
| POG lateral | .12 | .121 | .342 | .19 | .017 | .084 |
| 7-Up | .06 | .426 | .757 | .18 | .022 | .084 |
| 7-Down | .16 | .041 | .232 | .03 | .682 | .807 |
| PMQ-9 | .02 | .757 | .757 | .05 | .512 | .807 |
| PHQ-9 | .03 | .723 | .757 | –.02 | .830 | .876 |
| BAS Drive | .12 | .115 | .342 | .18 | .018 | .084 |
| BAS Reward | .04 | .652 | .757 | .09 | .251 | .477 |
| BIS | .11 | .160 | .389 | .03 | .722 | .807 |
| WASSUP | .12 | .115 | .342 | .15 | .060 | .162 |
| NGS | .05 | .515 | .757 | .04 | .605 | .807 |
| GSE | .07 | .368 | .757 | .04 | .591 | .807 |
| GAD-7 | .03 | .742 | .757 | .00 | .988 | .988 |
| PSWQ | .03 | .701 | .757 | .03 | .713 | .807 |
| SHAPS | .03 | .679 | .757 | .05 | .556 | .807 |
| IE-4 | –.04 | .602 | .757 | .09 | .238 | .477 |

### 3.4 Exploratory factor analysis

Our exploratory factor analysis revealed that at low resolution (*k* = 2–3), *µ* was most strongly correlated with a broad mania/approach factor loading on WASSUP (ambitious goal setting), POG (positive overgeneralization), NGS (narcissism), 7Up (hypomania), and BIS/BAS (behavioral activation) items (*r ≈.*15, Fig. 6B). At intermediate resolutions (*k* = 4–8), the strongest association shifted to a WASSUP-dominant factor (*r ≈.*15). While none of these correlations survived FDR correction applied within each level, the factor that was most strongly correlated with *µ* consistently loaded on mania-related items.

**Figure 6:**
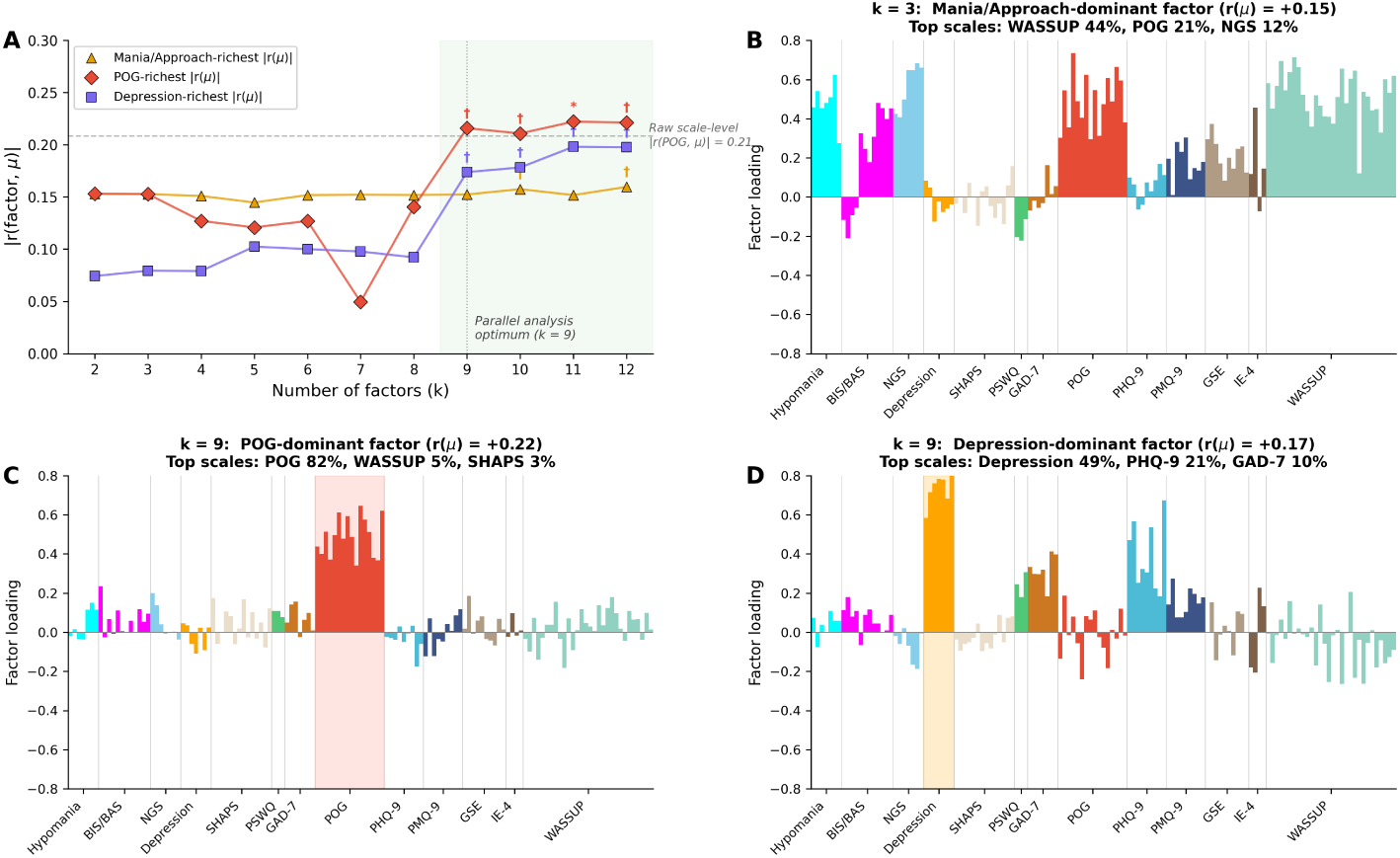
Exploratory factor analysis across levels of factor resolution. **A:** *|r*(factor*, µ*)*|* across factor solutions (*k* = 2–12) for the mania/approach-richest factor (orange triangles), the POG-richest factor (red diamonds), and the depression-richest factor (purple squares). Dashed line indicates the scale-level *|r*(POG*, µ*)*|* = .21; dotted line marks the parallel analysis optimum (*k* = 9). *^†^ p*_raw_ *<* .05; *^∗^ p*_FDR_ *<* .05. **B–D:** Factor loadings, with items grouped by scale along the *x*-axis. **B:** the factor most strongly correlated with *µ* at *k* = 3 (*r* = .15; WASSUP purity = 44%). **C:** the POG-dominant factor at *k* = 9 (*r* = .22; purity = 82%). **D:** the depression-dominant factor at *k* = 9 (*r* = .17; 7-Down purity = 49%), with the dominant scale’s items highlighted in each.

At high resolution (*k* = 9), POG items separated into a distinct factor, accounting for 82% of its loading variance (Fig. 6C). At this resolution, *µ* shifted to a specific POG-dominant factor (*r* = .22, *p*_raw_ = .006, *p*_FDR_ = .051), with the correlation remaining stable through *k* = 12 (*r* = .21–.22). A second factor, dominated by 7-Down items, also showed a modest association with *µ* (*r* = .17, *p*_raw_ = .026, *p*_FDR_ = .119; Fig. 6D). Notably, this depression-dominant factor was driven primarily by 7-Down items (purity = 49%), which assess trait-like depressive tendencies as part of a bipolar screening instrument, with secondary contributions from PHQ-9 items (21%) capturing current depressive symptoms. At the scale level, neither PHQ-9 nor GAD-7 showed an association with *µ*, suggesting that the behavioral signal relates to trait or episodic depressive content more than to current symptom severity. Beyond these two factors, only one additional factor showed *|r| > .*10 with *µ* at *k* = 9 — a WASSUP-dominant factor indexing ambitious goal-setting (*r* = .15, purity = 91%). All three factors with *|r| > .*10 thus loaded on bipolar-spectrum-relevant content (POG, trait depressive tendencies, and ambitious goal-setting), consistent with the broader interpretation that *µ* tracks individual differences along the bipolar spectrum. Loading profiles for all nine factors are shown in Fig. S5.

These results complement the scale-level confirmatory findings. In particular, the absence of association between *µ* and state measures of depression and anxiety is consistent across factor solutions, while the factor most strongly correlated with *µ* loads on mania-related items at every resolution and resolves most clearly onto positive overgeneralization when the factor structure affords sufficient separation of mania-adjacent constructs. At higher resolutions, a factor dominated by trait-like depressive items (7-Down) also showed an association with *µ*, suggesting that the behavioral signal may index a generalization mechanism implicated across bipolar spectrum traits, extending beyond positive overgeneralization to include trait-like depressive tendencies. Together, the item-level and scale-level results converge in suggesting that the Fossil Hunt task indexes a specific cognitive process — the breadth of reward generalization — that is linked to individual differences along the bipolar spectrum.

### 3.5 A computational mechanism of value generalization

Having established that measures of value generalization track self-reported POG and subclinical bipolar traits, we next asked what could modulate the breadth of generalization. From a reinforcement learning perspective, generalization allows value learned in one context to influence expectations in similar, unexperienced states (Gershman & Niv, 2015; Radulescu & Niv, 2019; Sutton & Barto, 2018; Wimmer et al., 2012; Wu et al., 2025). One key question is what might determine how broadly people generalize.

To answer this, we start from the assumption common in reinforcement learning that the value an agent assigns to a state depends, in part, on the best outcome it expects to reach from that state. For instance, an airport’s value depends not on the airport itself but on the best destination it lets you reach: a hub connecting to many desirable destinations is more valuable than one that does not, even though neither is a destination in itself. This forward-looking component allows value to propagate beyond directly experienced states, shaping expectations for situations the agent has not yet encountered.

Prior work has proposed that beliefs about self-efficacy modulate this forward projection, showing that low efficacy attenuates value propagation, resulting in avoidance behavior characteristic of anxiety (Zorowitz et al., 2020). In that framework, the value an agent anticipates from the next state is a weighted average of the best and worst actions available there, and the self-efficacy weight *w* sets their relative contribution. At *w* = 1 the agent anticipates the best available action, as in standard reinforcement learning; lower *w* gives more weight to the worst available action, so value decays more steeply across states and produces a narrower generalization gradient. We have previously proposed that a variant in which *w* itself increases without bound with success can give rise to behavior characteristic of hypomania (Li & Radulescu, 2024); the present simulation holds *w* fixed within each agent and varies it across agents.

In this model, action values *Q*(*s, a*) are updated via a reward prediction error (RPE):

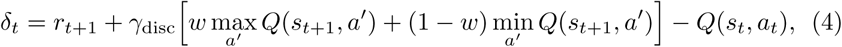

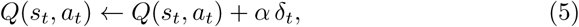

where *α ∈* (0, 1) is the learning rate, *r_t_*_+1_ is the experienced reward, and *γ*_disc_ *∈* (0, 1) is the temporal discount factor (we use *γ*_disc_ to avoid confusion with the lower asymptote *γ* in the psychometric model). The weight *w* takes values between 0 and 1, so standard maximizing Q-learning sets the upper bound on how broadly value can propagate in this model.

To test whether efficacy weighting is sufficient to shift the breadth of generalization, we simulated agents learning state values via temporal-difference learning (Sutton & Barto, 2018) on a linear track of 10 positions, moving one step left or right at each of the eight interior positions. The leftmost position terminated the episode with no reward; the rightmost terminated it with reward delivered on 85% of visits, matching the task’s reinforcement rate. We simulated 20 runs at each of five weights (*w ∈ {*0*, .*25*, .*50*, .*75, 1.00*}*), holding *α* = 0.1 and *γ*_disc_ = 0.8. Learned values were mapped to choice propensities via softmax and fit with the same four-parameter logistic procedure used for participant data.

Higher self-efficacy weights produced broader value gradients extending farther from the rewarded end of the track (Fig. 7A), and the corresponding choice gradients shifted accordingly (Fig. 7B). Mean simulated breadth increased monotonically across the five weights, from 15.4 at *w* = 0 to 46.8 at *w* = 1 (Fig. 7C). Because *w* = 1 recovers standard maximizing Q-learning, this shows that conservative weighting of future outcomes constrains how broadly value propagates, and that moving toward standard maximization yields correspondingly broader propagation. It does not produce breadth beyond what standard maximizing learning would produce. The simulated environment and response mapping also differ from the perceptual task, so simulated and behavioral breadth are analogous rather than equivalent quantities.

**Figure 7:**
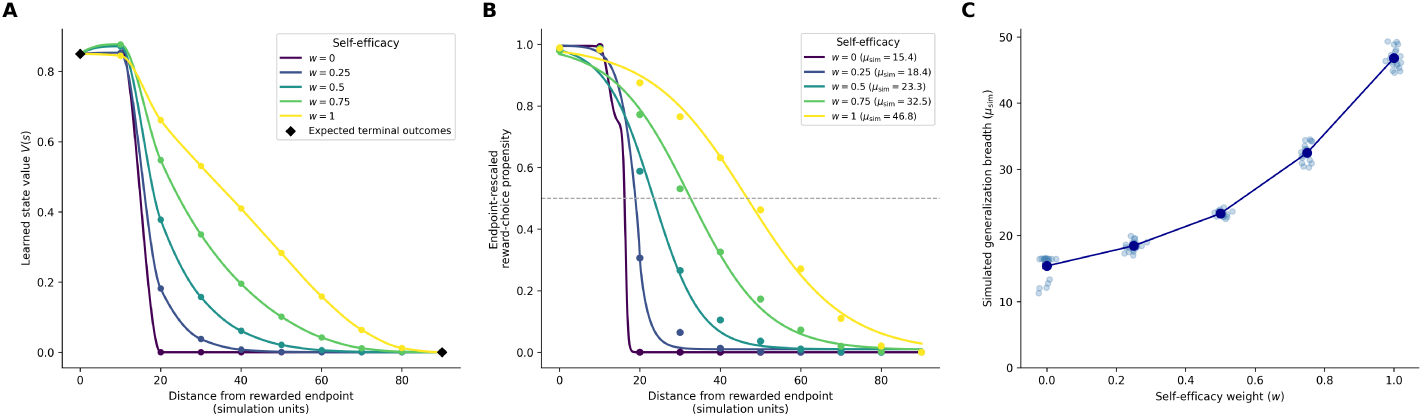
Simulated effect of self-efficacy weighting on reward generalization. At each next state, anticipated future value was *w* max*_a_′ Q*(*s^′^, a^′^*) + (1 *− w*) min*_a_′ Q*(*s^′^, a^′^*), with 0 *≤ w ≤* 1. **A:** Mean learned state values *V* (*s*) = max*_a_ Q*(*s, a*) at each weight; black diamonds mark the expected terminal outcomes. **B:** Mean choice gradients at each weight, with logistic fits and the corresponding *µ*_sim_ shown in the legend. **C:** Self-efficacy weight against fitted breadth.

## 4 Discussion

Positive overgeneralization (POG) refers to the tendency to generalize from specific successes to broad expectations of future reward, and has been linked to vulnerability to mania. Here we use a newly developed generalization task to show that this tendency corresponds to a measurable difference in how broadly learned value generalizes in an associative learning context. Consistent with our primary hypothesis, we found that individuals who scored higher in self-reported POG judged stimuli farther from the trained rewarded exemplar as valuable. This was reflected in the psychometric midpoint *µ*, which was positively associated with POG scores, and converged with a secondary response-time measure: higher POG was also associated with a shallower rise in response time as stimuli moved from the rewarded shape toward each participant’s own choice boundary. This pattern suggests that endorsing POG in self-report is accompanied by a measurable shift in how broadly learned value is applied to novel stimuli. We further found that this behavioral signal correlated with subclinical bipolar traits, suggesting it may be relevant to understanding vulnerability to bipolar disorder.

The effect size of the observed correlation between task-based generalization and POG was consistent with similar associations reported in computational psychiatry research linking behavioral task parameters to self-report symptom dimensions (Gillan, Kosinski, Whelan, Phelps, & Daw, 2016; Goldman et al., 2025). The response-time result adds process-level information about how readily each stimulus was judged valuable. Response times were fastest at the trained shapes and rose toward each participant’s choice boundary, peaking where a stimulus was equally likely to be judged valuable or not; moving from the rewarded shape toward that boundary, evidence for a “valuable” judgment weakens and responses slow accordingly. Higher-POG participants showed a shallower rise — stimuli partway toward the boundary were judged nearly as quickly as the rewarded shape itself — but were not slower overall: POG was unrelated to mean log RT, mean raw RT, and median raw RT.

These results suggest that positive overgeneralization is related to two separable aspects of the decision. The midpoint *µ* indexes where the boundary between valuable and non-valuable falls, while the distance slope *β_D_* indexes how strongly proximity to the trained shapes governs decision speed. That these two measures were themselves unrelated suggests they capture distinct consequences of the same cognitive style rather than two aspects of a single process.

The pattern of findings across confirmatory and exploratory analyses is also informative. POG was designated as the primary confirmatory construct given its direct theoretical alignment with the “Fossil Hunt” task: both operationalize the tendency to generalize positive outcomes, the former via self-report and the latter behaviorally. That both task measures showed their largest association with POG — while exploratory correlations with approach motivation measures did not survive correction for multiple comparisons — is therefore consistent with our hypotheses. The second confirmatory association, between *µ* and 7Up/7Down total, was also significant but smaller. Whether the non-significant exploratory associations with subscale scores reflect genuine null effects or true but underpowered relationships remains an open question that larger samples can resolve.

Beyond the scale-level analyses, exploratory factor analyses on item-level self-report data converged with the scale-level findings: across factor resolutions, *µ* tracked factors loading on mania-related items, loading most cleanly onto a POG-dominant factor at the parallel analysis optimum. A secondary association with a trait-depression factor emerged at higher resolutions, while state measures of depression and anxiety remained unassociated with *µ* throughout. This trait-state dissociation is theoretically informative: it suggests that broader reward generalization reflects a stable cognitive style rather than a consequence of current mood, consistent with POG as a vulnerability factor rather than a symptom state.

### 4.1 Theoretical Implications

We propose self-efficacy-weighted value propagation as a candidate computational mechanism underlying individual differences in generalization breadth. In this model, self-efficacy sets how much an agent anticipates the best rather than the worst outcome available from the next state. Although the model uses a sequential state space, whereas the task here involves judgments across a perceptual continuum, both instantiate the same abstract pattern: value is assigned to states or stimuli increasingly distant from a reinforced reference point. The simulation results establish that efficacy weighting is sufficient to modulate generalization breadth. A dynamic extension in which self-efficacy updates with experience (Li & Radulescu, 2024) offers one route by which success could move an initially conservative agent toward broader generalization of attainable outcomes.

### 4.2 Clinical Implications and Future Directions

POG has been identified as a risk-relevant cognitive style in hypomania and mania (Stange et al., 2012), but has until now been assessed primarily through self-report. The present findings have two potential clinical implications. First, if positive overgeneralization is a stable individual difference, it could serve as a behavioral marker for identifying individuals at elevated risk for mania, independently of the insight required by self-report measures. Second, interventions targeting the cognitive processes underlying overgeneralization – such as self-efficacy calibration – could be evaluated using task-based measures, offering a behavioral index of treatment response that complements self-report. A particularly important next step will be to examine whether task-based measures track mood state dynamically (for instance, do individuals show broader generalization during hypomanic episodes compared to euthymic baselines?), which would strengthen its potential utility as a clinical monitoring tool.

### 4.3 Limitations

Several limitations warrant consideration. First, the sample was recruited online and may not generalize to clinical populations; testing whether task-based measures of positive overgeneralization predict mania risk in individuals with bipolar disorder or those at clinical high-risk remains an important next step.

Second, the cross-sectional design precludes causal inference – we cannot determine whether broader generalization precedes or follows the development of POG as a cognitive style. Longitudinal designs tracking POG alongside mood episodes would be informative.

Third, the simulation is a qualitative proof of principle rather than a model fit to participant choices. Its sequential environment and mapping from learned values to choices are simplifying assumptions, and other mechanisms could produce similar gradients. Pinpointing the exact mechanism requires paradigms designed to elicit self-efficacy beliefs directly. Linking such estimates to positive overgeneralization at the individual level, and testing whether self-efficacy mediates the association between POG and symptoms, is an important direction for future work.

Finally, the association with bipolar-spectrum traits was carried by different components for the two behavioral measures — 7-Down for *µ* and 7-Up for *β_D_*— and neither measure related to current manic symptoms. The behavioral signal therefore appears to track trait-like content across the bipolar spectrum rather than mania specifically, and its relationship to mania risk remains to be established in samples with greater symptom range.

### 4.4 Conclusion

This work introduces a behavioral and computational framework for studying positive overgeneralization – a cognitive style linked to mania vulnerability that has until now not been precisely quantified. By demonstrating that a psychometric parameter derived from a reward generalization task tracks both self-reported POG and subclinical bipolar traits, we provide an objective behavioral measure on a construct previously accessible only through self-report. The proposed self-efficacy mechanism suggests that how much people anticipate the best available outcome may set how broadly value propagates, offering one candidate explanation for the individual differences observed in excessive goal pursuit characteristic of mania.

## Supplementary Materials

### S1 Stimulus generation

Stimuli were generated following the procedure developed by Malaviya et al. (2022). Each stick-figure shape was defined by 9 continuous features describing the configuration of the body and limbs. Two endpoint stick figures were selected to yield maximally distinct shapes, and eight intermediate stimuli were generated by linearly interpolating feature values between the two endpoints, yielding ten total stimulus positions along the shape dimension (Fig. S1). For the present study, the stick-figure quadrupeds were re-labeled as “dinosaur fossils” to fit the cover story of the Fossil Hunt task, and reward assignment to the two endpoint shapes was counterbalanced across participants.

### S2 Perceptual validation of the stimulus continuum

A separate sample of 49 participants rated the pairwise similarity of the ten stimulus shapes on a 0–4 scale. Mean rated similarity decreased monotonically with distance along the shape dimension (Fig. S2A). Non-metric multidimensional scaling recovered the stimulus order exactly (Spearman *ρ* = 1.00), with recovered coordinates falling close to even spacing (Fig. S2C), and distance along the shape dimension accounted for 98% of the variance in mean pairwise dissimilarity. The stimuli therefore lie along a single perceptual dimension at roughly equal intervals.

### S3 Psychometric function fitting details

Parameters were estimated via nonlinear least-squares using scipy.optimize. curve fit with the trust-region-reflective algorithm. For each participant, binary reward predictions were averaged within the 10 stimulus bins on the distance-from-reward coordinate *d*, yielding a proportion “valuable” response at each position. Initial values were *µ*_0_ = 45 (median stimulus value), *σ*_0_ = 1, *γ*_0_ = min(*y*_obs_), *λ*_0_ = max(*y*_obs_), with bounds *µ ∈* [0, 90], *σ ∈* (0*, ∞*), *γ ∈* [0, 1], *λ ∈* [0, 1]. Maximum function evaluations were set to 20,000. Each participant completed 100 test trials (10 per stimulus bin).

### S4 Parameter recovery

To assess whether the fitting procedure could reliably recover individual differences, we conducted a parameter recovery simulation. For each of the 163 participants, we generated 200 synthetic datasets by simulating binary responses from their fitted four-parameter logistic model using the same stimulus values and trial counts as the actual experiment. Each synthetic dataset was then refit using the identical estimation procedure (same optimizer, bounds, and initial values; random seed = 42). Recovery was assessed by correlating the generating (true) parameter values with the mean recovered values across simulations (Fig. S3A). Recovery was excellent for all parameters: *µ* (*r* = .988), *σ* (*r* = .920), *γ* (*r* = .971), *λ* (*r* = .918), and AUC (*r* = .998). The transition scale *σ* showed a modest negative bias (mean bias = *−*1.10), likely reflecting compression toward the lower bound for participants with steep psychometric functions; all other parameters showed negligible bias (*|*mean bias*| <* .02). Full recovery results are shown in Fig. S3A.

Recovery of the RT coefficients was assessed with a separate two-stage simulation. For each participant, 200 datasets were generated: choices were simulated from their fitted psychometric function and refit, log RT was generated from their fitted RT model, and distance was recomputed from the *recovered* boundary before both coefficients were re-estimated. Across individual task-length datasets, median true–recovered correlations were .82 for *β_D_* (central 95% simulation interval [.77, .86]) and .80 for *β_A_* ([.71, .84]). Holding the generating midpoint fixed left *β_D_* recovery essentially unchanged (median *r* = .82) but improved *β_A_* (median *r* = .84) and brought its interval coverage from .90 to .95, indicating that uncertainty in the fitted boundary propagates mainly into the directional coefficient.

### S5 Parameter intercorrelations

We examined pairwise correlations among the fitted parameters to determine whether they capture distinct aspects of behavior (Fig. S3B). As expected, *µ* and AUC were strongly positively correlated (*r* = .881, *p <* .001), since AUC is computed from the same fitted curve; we therefore report AUC descriptively rather than as independent evidence. The midpoint and transition scale were unrelated (*r* = *−*.036, *p* = .651). The lower and upper asymptotes were moderately negatively correlated (*r* = *−*.291, *p <* .001), and *γ* was modestly positively associated with AUC (*r* = .265).

The matrix also includes the jointly estimated RT coefficients. Generalization breadth was unrelated to the distance slope *β_D_* (*r* = .027, *p* = .732) but positively associated with the directional asymmetry coefficient *β_A_* (*r* = .346, *p <* .001); *β_D_* and *β_A_* were modestly negatively correlated (*r* = *−*.162, *p* = .039).

### S6 Factor loadings at the parallel-analysis optimum

To complement the factor analysis results reported in the main text (Fig. 6), Fig. S5 shows loading profiles for all nine factors extracted at the parallel analysis optimum (*k* = 9), ordered by *|r*(*µ*)*|*. The three factors associated with *µ* at *|r| >* .10 were POG-dominant (purity = 82%, *r* = .22), depression-dominant (primarily 7-Down; purity = 49%, *r* = .17), and WASSUP-dominant (purity = 91%, *r* = .15). The remaining six factors captured hypomania, BIS/BAS, GSE, anxiety/worry, SHAPS, and NGS dimensions, none correlating with *µ* above *|r|* = .10.

### S7 Psychometric response steepness

The fitted *σ* parameter describes the width of the psychometric transition rather than its slope. We therefore quantified maximum steepness as

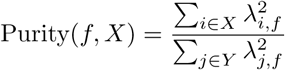

the maximum derivative of the fitted function, attained at *d* = *µ*. The raw distribution was strongly right-skewed (skewness = 3.33), so the individual-difference metric was log *S*, which reduced skewness to 1.26 (Fig. S7A–B).

Twenty-four participants had transition scales below one stimulus unit, well under the 10-unit spacing of the presented stimuli; these appear as a separate cluster at the upper end of the log *S* distribution (Fig. S7B). For these participants *σ* is constrained by the resolution of the design rather than estimated from the data. Because the task presents stimuli at 10-unit intervals, it cannot resolve transitions narrower than one step, and log *S* is therefore not a well-identified individual-difference measure in this design. For a substantial minority of participants it reflects the spacing of the stimulus set rather than the sharpness of their psychometric function. We therefore report the distribution of steepness for completeness but do not report or interpret its correlations with self-report.

### S8 Response-time distributions

Raw RTs were right-skewed at every stimulus position (skewness range 2.35– 4.98); natural-log transformation reduced this range to .60–1.34 (Fig. S8). Distributions near the middle of the stimulus range were shifted toward longer RTs, consistent with the boundary-centered slowing reported in the main text.

### S9 Log-RT asymmetry and rewarded-direction slope

POG total showed a modest positive correlation with reward-positive *β_A_* (*r* = .19, *p* = .014, *q* = .162; Fig. S9A) that did not survive FDR correction across all 19 questionnaire scores; no score survived correction for *β_A_*.

We defined *β*^+^ = *β_D_* + *β_A_* as the rewarded-direction log-RT distance slope. POG total was positively associated with *β*^+^ (*r* = .30, *p <* .001, *q* = .002). The upward (*r* = .23, *q* = .014), social (*r* = .28, *q* = .003), and lateral (*r* = .27, *q* = .004) subscales also survived FDR correction across all 19 scores; no non-POG score survived correction (Fig. S9B; Table S2).

### Ethics statement

The Program for the Protection of Human Subjects at the Icahn School of Medicine at Mount Sinai determined this study to be exempt from IRB review (study no. IF2813888, protocol “Online Studies of Naturalistic Reinforcement Learning in Psychiatric Disorders”). Participants were recruited anonymously via Prolific and provided informed consent by reading a research information sheet and clicking to indicate agreement before beginning the study; they could withdraw at any time by closing the browser. The study posed minimal risk and collected no identifiable information.

---

**Figure S1:**
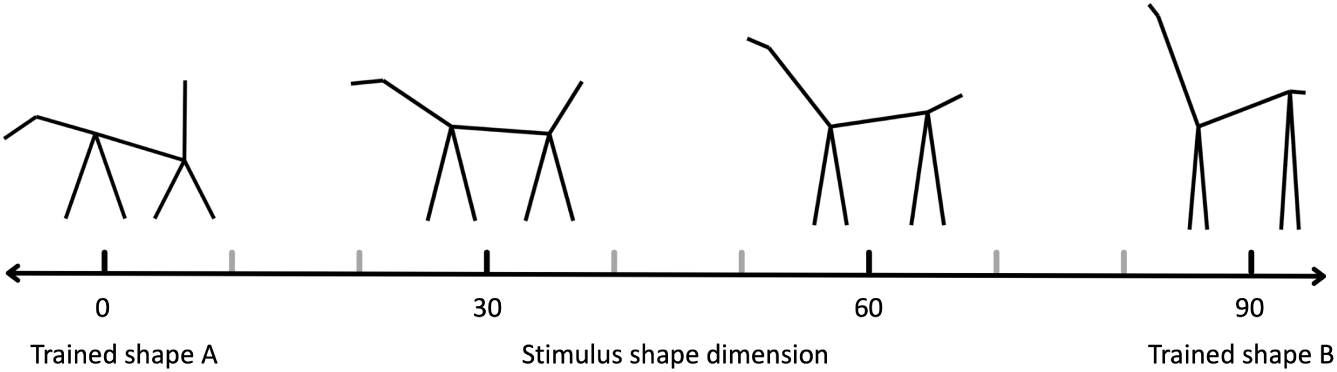
Examples of stimuli along the shape dimension used in the Fossil Hunt task. Four representative stick-figure stimuli at evenly spaced positions (0, 30, 60, 90) along the continuous shape dimension. Black tick marks indicate the four positions shown above the axis; gray tick marks indicate the six additional positions (10, 20, 40, 50, 70, 80) presented during the task but omitted from the figure for clarity, yielding ten stimulus positions in total. The two endpoint shapes (positions 0 and 90, labeled “Trained shape A” and “Trained shape B”) were used during the training phase; the eight intermediate shapes were algorithmically generated by linearly interpolating feature values between the endpoints across nine continuous shape features. Reward assignment to the two endpoint shapes was counterbalanced across participants.

**Figure S2:**
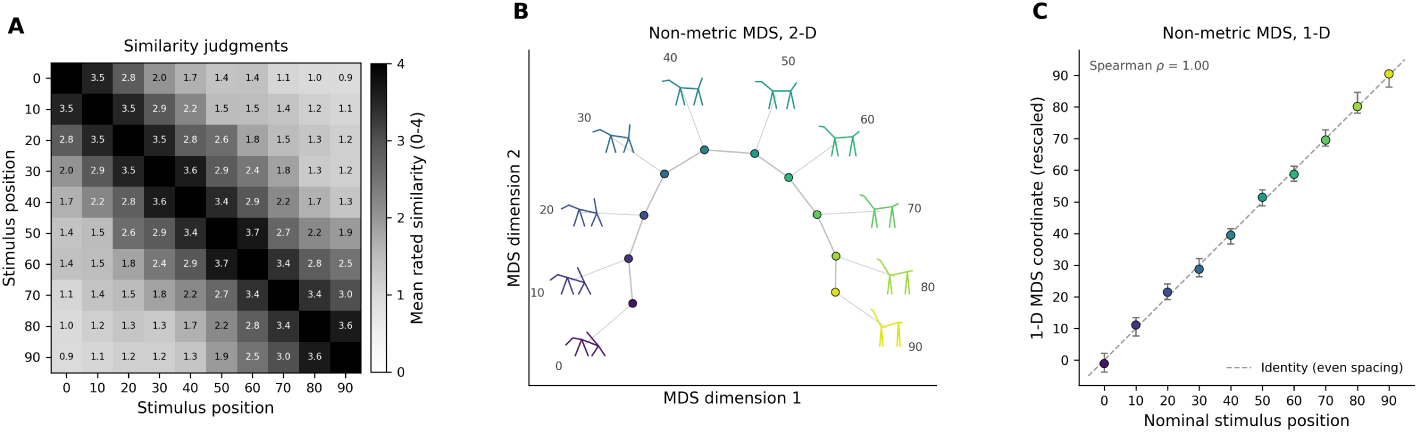
Perceptual validation of the stimulus continuum. **A:** Mean pairwise similarity ratings between the ten stimulus positions. **B:** Two-dimensional MDS solution, with the corresponding stick figures shown; the arc is the curvature typical of a single underlying dimension. **C:** One-dimensional MDS coordinates against nominal stimulus position. Error bars are 95% participant bootstrap intervals

**Figure S3:**
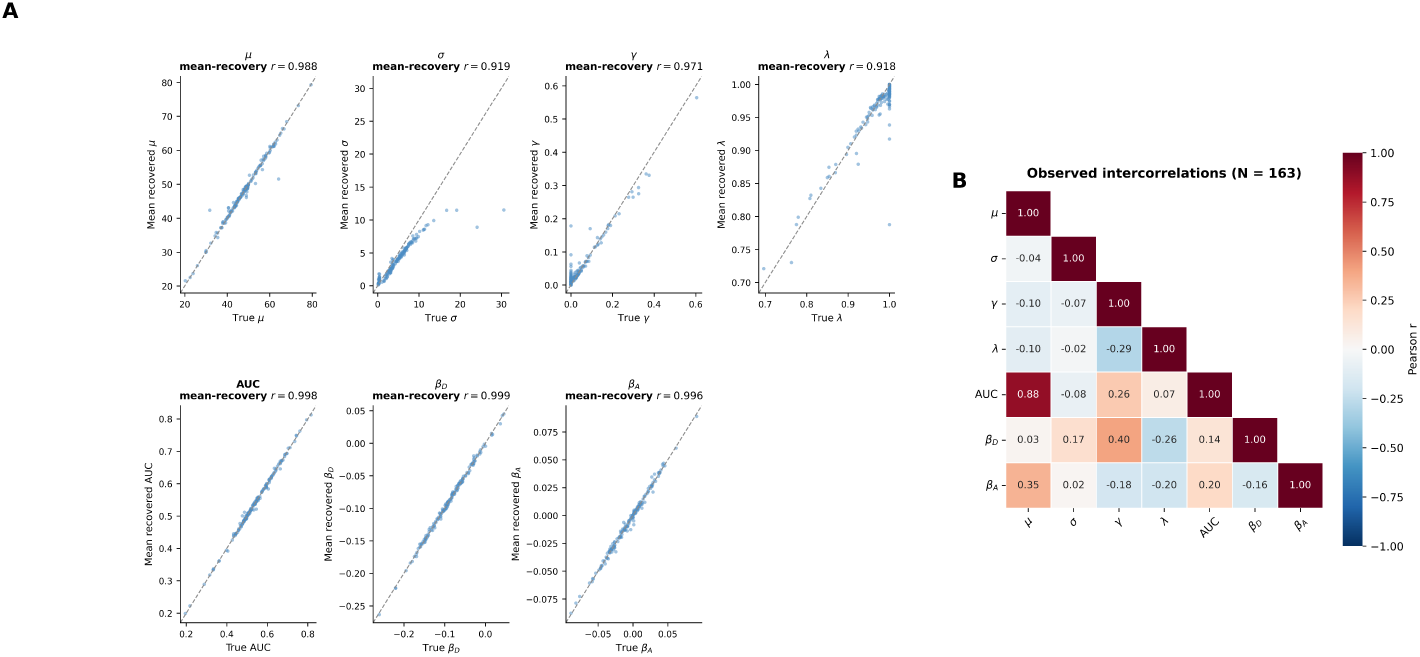
Parameter recovery and descriptive intercorrelations. **A:** True parameter values against participant-mean recovered values across 200 synthetic datasets; dashed lines denote identity. Facet annotations give the correlation between true and mean recovered estimates. For *β_D_* and *β_A_*, choices and log RT were simulated using the observed trial structure, the psychometric boundary was refit, and distance was recomputed before refitting the RT model. Because each point averages 200 datasets, these correlations reflect mean-estimate fidelity rather than single-dataset precision; replicate-wise recovery is reported in the text. **B:** Pairwise Pearson correlations among fitted parameters (*N* = 163). *β_A_* uses the signed coordinate, positive toward the rewarded endpoint.

**Figure S4:**
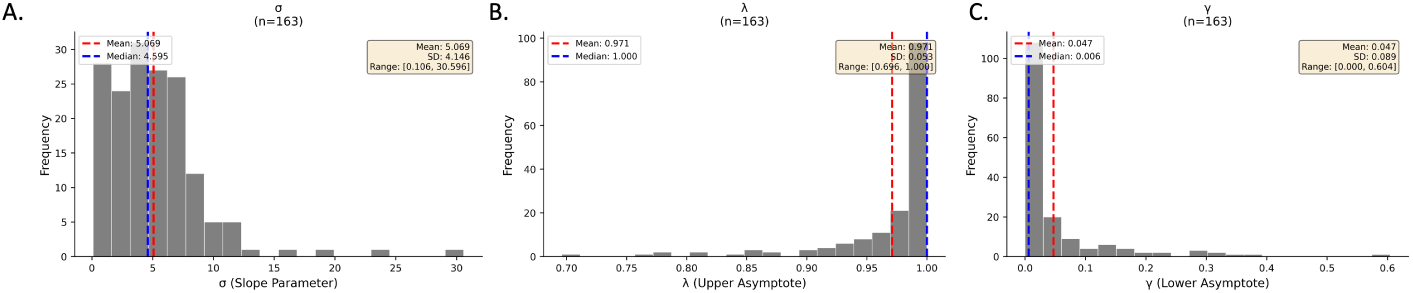
Distributions of additional fitted parameters. **A:** Transition scale *σ* (*M* = 5.07, *SD* = 4.15), showing a right-skewed distribution reflecting a subset of participants with steep, near-step-function transitions. **B:** Upper asymptote *λ* (*M* = .971, *SD* = .053), with most participants near ceiling. **C:** Lower asymptote *γ* (*M* = .047, *SD* = .089), with most participants near floor. Dashed red and blue lines indicate mean and median, respectively (*N* = 163). The distribution of *µ* is shown in the main text (Fig. 2C).

**Figure S5:**
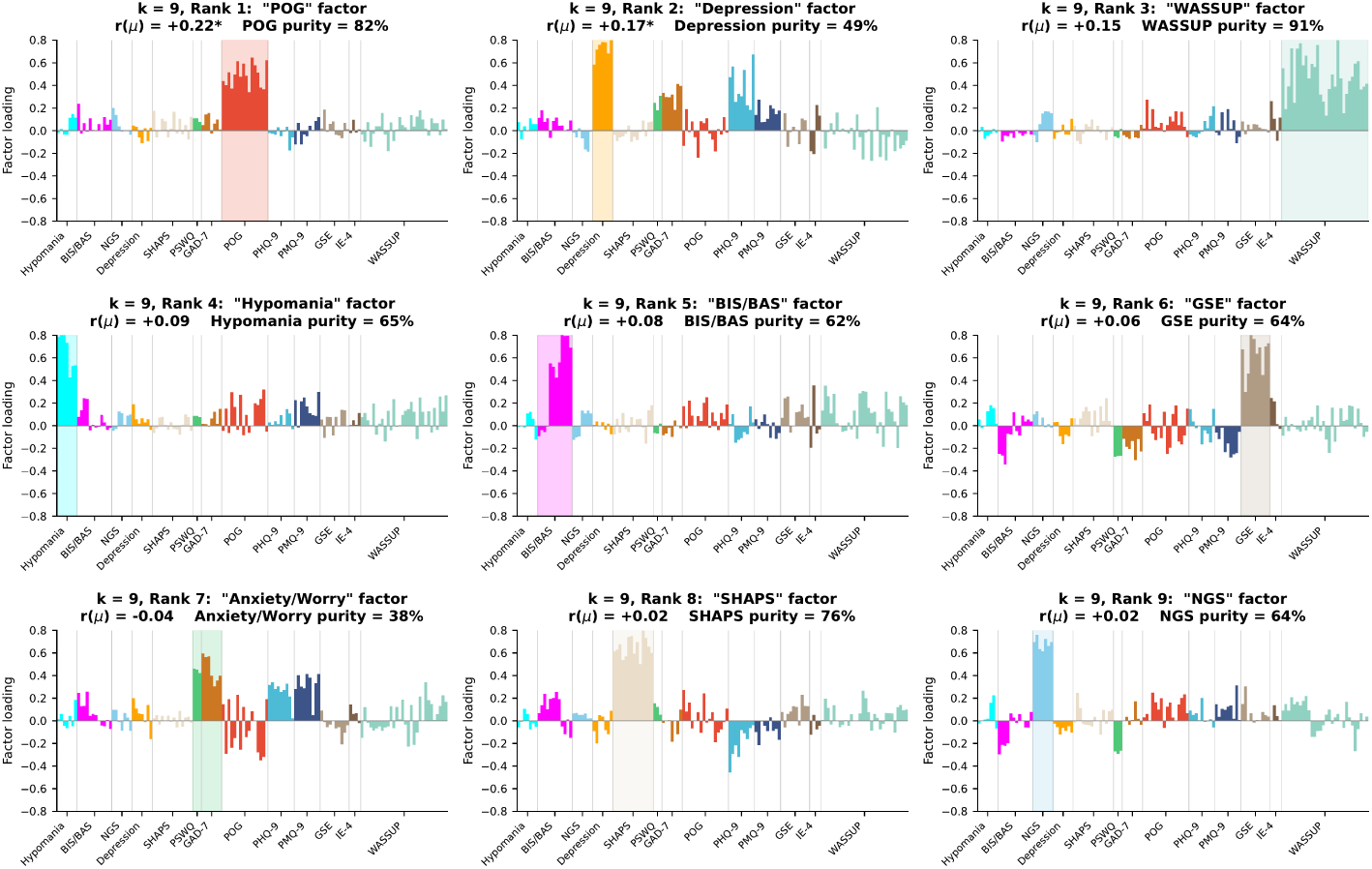
Loading profiles of all nine factors at the parallel analysis optimum (*k* = 9). Each panel shows the factor loadings for one of the nine factors extracted at *k* = 9, ordered by *|r*(*µ*)*|* descending). Items are grouped along the *x*-axis by self-report scale. Each factor is labeled by the scale contributing the largest share of its squared loadings, with that scale’s items highlighted within the panel. Annotations show the Pearson correlation between factor scores and *µ* (*r*(*µ*)) and the dominant scale’s purity, defined as the proportion of the factor’s total squared loadings contributed by items from that scale. *N* = 163. *^†^* raw *p <* .05; *^∗^* FDR-corrected *p <* .05.

**Figure S6:**
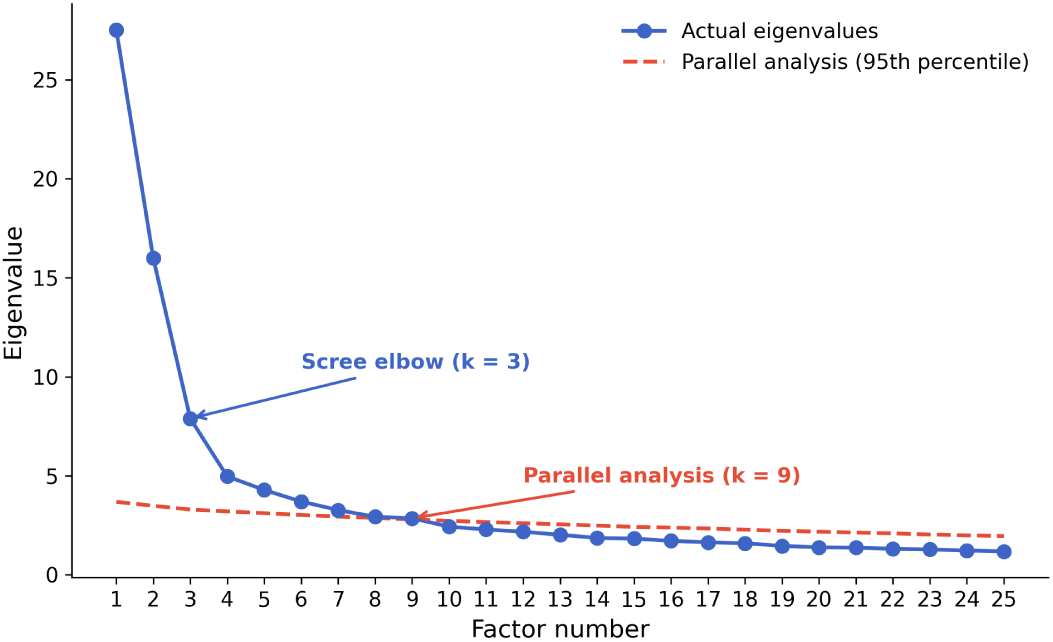
Factor selection criteria. Scree plot of eigenvalues from the observed correlation matrix (solid blue) and 95th percentile eigenvalues from parallel analysis with 1000 random permutations (dashed red). The scree elbow suggests *k* = 3; parallel analysis indicates *k* = 9 factors with eigenvalues exceeding the random threshold.

**Figure S7:**
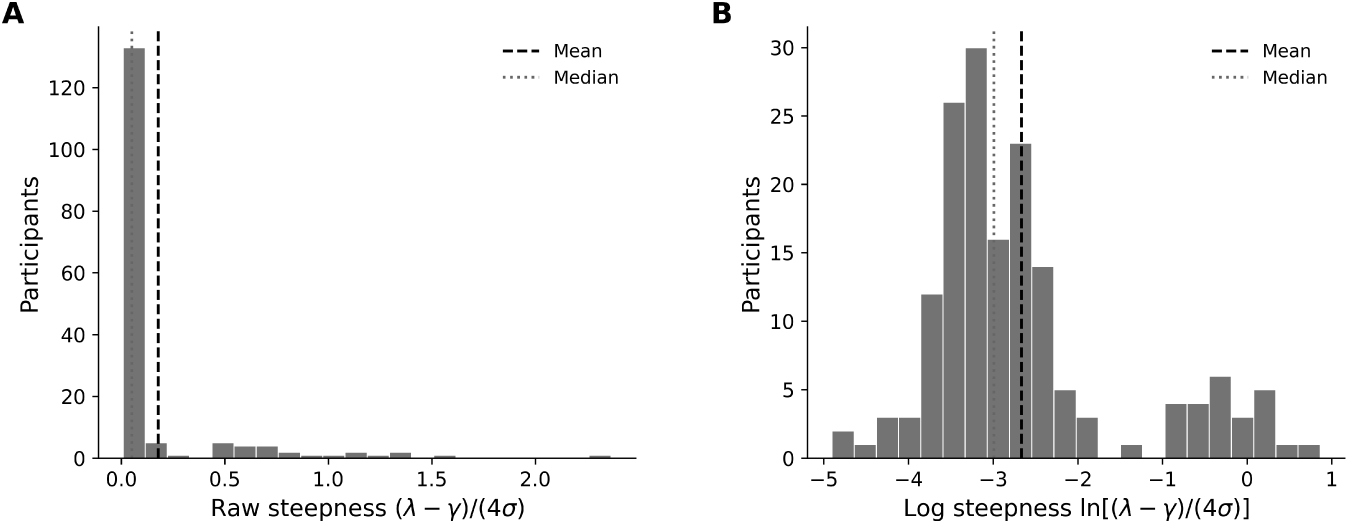
Psychometric response steepness. **A:** Distribution of the raw maximum derivative (*λ − γ*)*/*(4*σ*). **B:** Distribution after natural-log transformation. contributions near the middle of the stimulus range were shifted toward longer RTs, consistent with the boundary-centered slowing reported in the main text.

**Figure S8:**
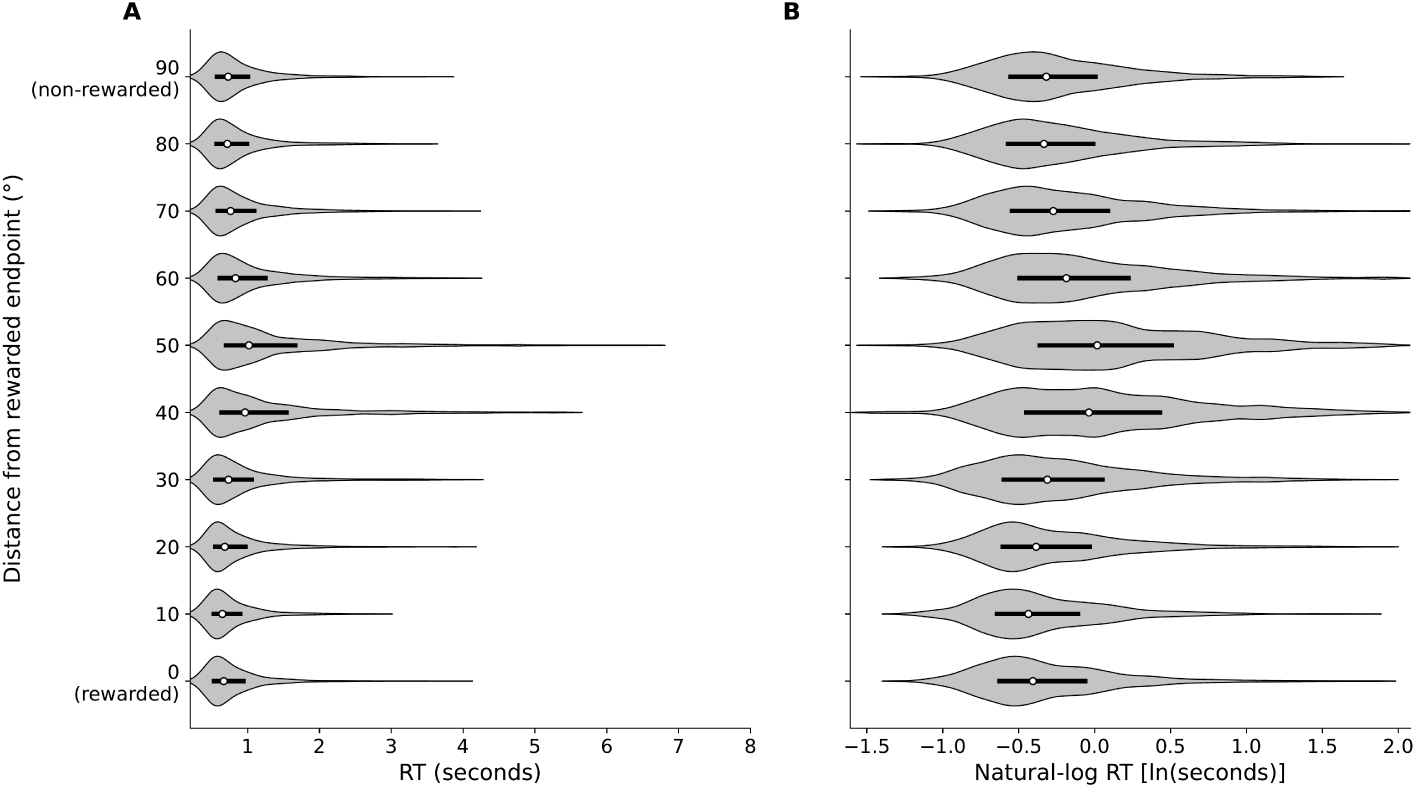
Response-time distributions by distance from the rewarded endpoint. **A:** Raw test-trial RT distributions for the ten stimulus positions. **B:** Distributions of the same trials after natural-log transformation. The reporting coordinate is 0 at the trained rewarded endpoint and 90 at the trained nonrewarded endpoint.

**Figure S9:**
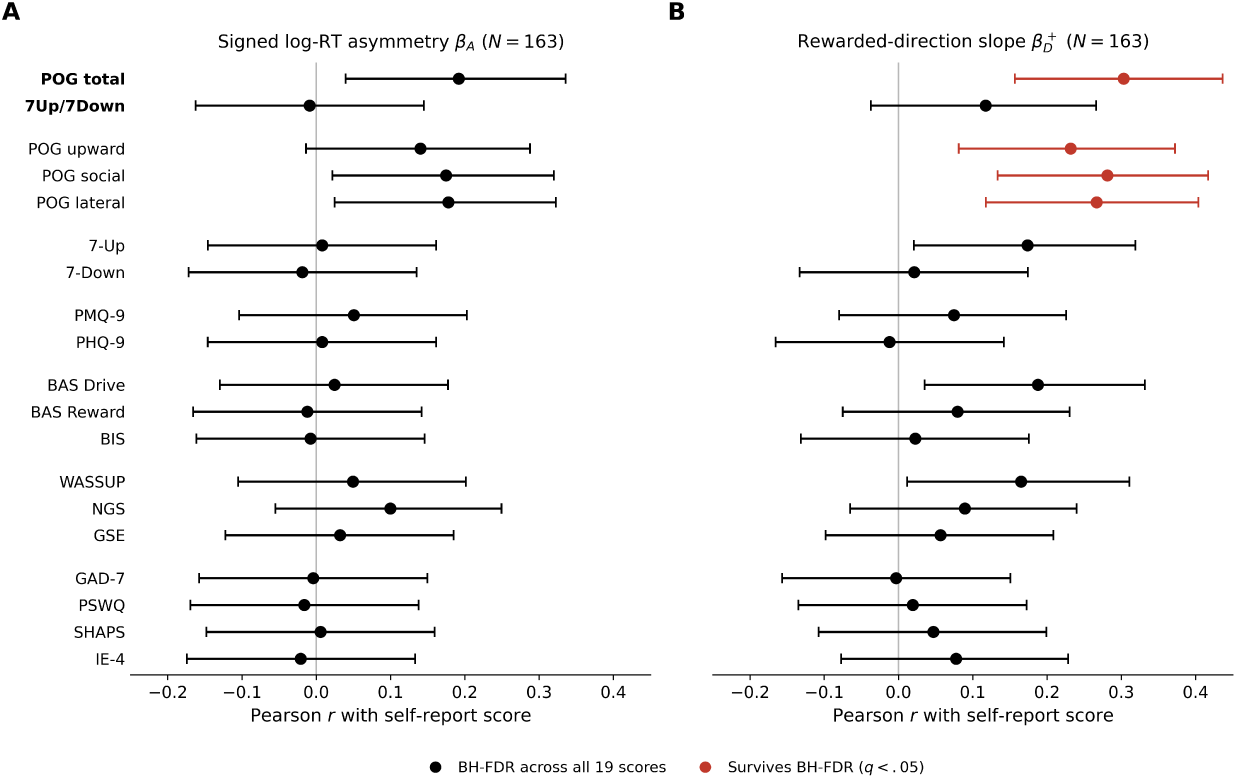
Self-report correlates of the log-RT asymmetry *β_A_* and the rewarded-direction slope *β*^+^. Pearson correlations with 95% confidence intervals between each of the 19 self-report scores and subscales and **(A)** the signed log-RT asymmetry coefficient *β_A_* and **(B)** the rewarded-direction slope *β*^+^ = *β_D_* + *β_A_* (*N* = 163). FDR was applied across all 19 scores within each panel. Red markers denote correlations surviving correction at *q <* .05 (POG total and the three POG subscales in panel B; none in panel A).

**Table S1:** Pearson correlations between *µ* and factor scores at *k* = 9 (*N* = 163). Each factor is labeled by the scale contributing the largest share of its squared loadings. Purity indicates the proportion of the factor’s total squared loadings contributed by items from the dominant scale. FDR correction was applied across the nine factor–*µ* correlations at this resolution.

| Rank | Factor (dominant scale) | $r$ | 95% CI | $p_{\text{raw}}$ | $p_{\text{FDR}}$ | Purity |
| --- | --- | --- | --- | --- | --- | --- |
| 1 | POG | .22 | [.06, .36] | .006 | .051 | .82 |
| 2 | Depression (7Down) | .17 | [.02, .32] | .026 | .119 | .49 |
| 3 | WASSUP | .15 | [−.00, .30] | .052 | .156 | .91 |
| 4 | Hypomania | .09 | [−.07, .24] | .277 | .605 | .65 |
| 5 | BIS/BAS | .08 | [−.08, .23] | .336 | .605 | .62 |
| 6 | GSE | .06 | [−.09, .21] | .432 | .648 | .64 |
| 7 | Anxiety/Worry | −.04 | [−.20, .11] | .583 | .749 | .38 |
| 8 | SHAPS | .02 | [−.14, .17] | .819 | .841 | .76 |
| 9 | NGS | .02 | [−.14, .17] | .841 | .841 | .64 |

**Table S2:** Secondary Pearson correlations of all 19 questionnaire scores with the reward-positive log-RT asymmetry *β_A_* and rewarded-direction slope *β*^+^ = *β_D_* + *β_A_* (*N* = 163). All *p*-values are unadjusted and two-sided; *q*-values apply separately across all 19 scores for each behavioral measure, including POG total and the 7Up/7Down composite.

| Score | $\beta_A$ | | | $\beta_D^+$ | | |
| --- | --- | --- | --- | --- | --- | --- |
| | $r$ | $p_{\text{raw}}$ | $q$ | $r$ | $p_{\text{raw}}$ | $q$ |
| POG total | .19 | .014 | .162 | .30 | < .001 | .002 |
| 7Up/7Down | -.01 | .909 | .959 | .12 | .136 | .323 |
| POG upward | .14 | .074 | .351 | .23 | .003 | .014 |
| POG social | .17 | .026 | .162 | .28 | < .001 | .003 |
| POG lateral | .18 | .023 | .162 | .27 | < .001 | .004 |
| 7-Up | .01 | .919 | .959 | .17 | .027 | .084 |
| 7-Down | -.02 | .812 | .959 | .02 | .789 | .902 |
| BIS | -.01 | .921 | .959 | .02 | .774 | .902 |
| BAS Drive | .02 | .756 | .959 | .19 | .016 | .062 |
| BAS Reward | -.01 | .878 | .959 | .08 | .313 | .544 |
| PMQ-9 | .05 | .521 | .959 | .07 | .343 | .544 |
| PHQ-9 | .01 | .920 | .959 | -.01 | .879 | .928 |
| WASSUP | .05 | .531 | .959 | .17 | .035 | .095 |
| NGS | .10 | .206 | .781 | .09 | .257 | .542 |
| GSE | .03 | .684 | .959 | .06 | .474 | .693 |
| GAD-7 | .00 | .959 | .959 | .00 | .968 | .968 |
| PSWQ | -.02 | .837 | .959 | .02 | .807 | .902 |
| SHAPS | .01 | .942 | .959 | .05 | .552 | .750 |
| IE-4 | -.02 | .791 | .959 | .08 | .326 | .544 |

